# *Toxoplasma* controls host cyclin E expression through the use of a novel MYR1-dependent effector protein, HCE1

**DOI:** 10.1101/573618

**Authors:** Michael W. Panas, Adit Naor, Alicja M. Cygan, John C. Boothroyd

## Abstract

*Toxoplasma gondii* is an obligate intracellular parasite that establishes a favorable environment in the host cells in which it replicates. We have previously reported that it uses MYR-dependent translocation of dense granule proteins to elicit a key set of host responses related to the cell cycle, specifically E2F transcription factor targets including cyclin E. We report here the identification of a novel *Toxoplasma* effector protein that is exported from the parasitophorous vacuole in a MYR1-dependent manner and localizes to the host’s nucleus. Parasites lacking this inducer of <u>H</u>ost <u>C</u>yclin <u>E</u> (HCE1) are unable to modulate E2F transcription factor target genes and exhibit a substantial growth defect. Immunoprecipitation of HCE1 from infected host cells shows that HCE1 efficiently binds elements of the cyclin E regulatory complex, DP1 and its partners E2F3 and E2F4. Expression of HCE1 in *Neospora caninum*, or in uninfected HFFs, shows localization of the expressed protein to the host nuclei and strong cyclin E up-regulation. Thus, HCE1 is a novel effector protein that is necessary and sufficient to impact the E2F-axis of transcription resulting in co-opting of host functions to *Toxoplasma’s* advantage.

## Introduction

Intracellular infectious agents face unique challenges and opportunities. One such is interfacing with the host cell cycle, and many have evolved ways to speed up, slow down, or otherwise disrupt this process. *Toxoplasma gondii* is an obligate intracellular eukaryote that conforms to this rule. Indeed, this ubiquitous member of the phylum *Apicomplexa* has previously been described to cause host cells to stall in states ranging between S phase and the G2/M transition (1). In some host cells, this manifests in the host cell endocycling, duplicating its DNA without subsequent cytokinesis, and previous evidence has suggested that this is mediated by an active, but unknown, parasite-derived activity (2).

*Toxoplasma* tachyzoites use secreted effectors derived from the dense granules to manipulate host cell functions while replicating in the parasitophorous vacuole (PV)(3, 4). A select set of these dense granule proteins can cross the parasitophorous vacuole membrane (PVM) and enter the host cytoplasm (5) in a process that is dependent on at least four parasite proteins; three of these are located at the PVM and have been termed MYR1, MYR2, and MYR3 (2, 6) while a fourth, aspartyl protease 5 (ASP5), is found within the Golgi and catalyzes proteolysis at a conserved Arg-Arg-Leu (RRL) sequence (7, 8). To date, GRA16 (6, 9), GRA24 (2) and GRA18 (10) have been shown to employ this machinery and the loss of the TgIST-induced phenotype in Myr^−^ mutants is consistent with it being a fourth such protein (11).

Using mutants defective in MYR1 and ASP5, and human foreskin fibroblasts (HFFs) as the host cell, we recently described the totality of impacts on the host transcriptome that are dependent on effectors that use this system to translocate across the PVM. The data showed the expected impacts on host processes already known to be caused by GRA16, GRA24, GRA18, and TgIST. They also showed, however, a profound and unexplained MYR1-dependent impact on gene sets defined as E2F targets and/or G2/M checkpoint control (12). The E2F transcription factors are a family of DNA binding proteins that form a heterodimer with Dimerization Partner 1 (DP1) and regulate a cohort of genes including cyclin E and its cyclin dependent kinase (CDK2) which control progression of the cell cycle (13).

To identify the effector responsible for these E2F-mediated effects, we took a bioinformatic approach, focusing on candidate proteins that would be predicted to traffic across the PVM, reach the host nucleus and impact the host cell cycle. This approach proved successful and we describe here a novel *Toxoplasma* protein whose entry into the host cell is MYR1-dependent, that binds to E2F/DP1 heterodimers and is both necessary and sufficient for upregulation of host cyclin E, a key regulator of the host cell cycle.

## Results

### TGGT1_239010 Is a Dense Granule Protein that Traffics to the Host Nucleus in a MYR1-and ASP5-Dependent Manner

Similarities exist between the four known effector proteins that transit across the PVM and we sought to harness this information to identify the effector that might be mediating the up-regulation of cyclin E. Specifically, the known effectors all originate in the dense granules, are exported from the parasite, and do not end up inserted in a membrane but rather transit to the host cytosol or nucleus where many then drive a function that is specific to cells infected with *Toxoplasma* compared to those infected with the closely related genus, *Neospora caninum*. The latter point was specifically true in the case of cyclin E where transcriptomic analyses have showed that *Toxoplasma* upregulates this gene strongly while the data for *Neospora* revealed no such impact (14).

Based on all this, we searched existing excreted secreted antigen lists (15) for proteins predicted to have: 1) a signal sequence for export; 2) no transmembrane domain that might prevent translocation across the PVM; 3) a nuclear localization signal to mediate import into the host nucleus; and 4) either no orthologue in *Neospora caninum* or a very low level of similarity with such a gene. For the first two criteria, we used the predictions in ToxoDB (version 27, released Feb 19, 2016) and then confirmed these with the publically available prediction software Phobius (http://phobius.sbc.su.se/), a combined transmembrane topology and signal peptide predictor.

One gene that met these criteria was *TGGT1_239010*, which is predicted to encode a 685 amino acid protein with a signal sequence and no transmembrane domains (Fig. 1A). Previously published phosphoproteomic data examining differential phosphorylation states of proteins in parasites within intact vacuoles vs. syringe-released parasites gives insight into whether these post translational modifications are done inside the parasite or within the host/PV. While this data set cannot distinguish between Host/PV, it indicates that TGGT1_239010 is phosphorylated at 4 serines after being secreted from the parasite (ToxoDB, (16)), consistent with phosphorylation of other effector proteins. TGGT1_239010 also contains a predicted monopartite NLS downstream of the signal sequence that was identified by NLS mapper (http://nls-mapper.iab.keio.ac.jp/cgi-bin/NLS_Mapper_form.cgi) as AHRKKRRQL with a score of 8.5, suggesting a high likelihood of being a eukaryotic NLS (7 is the threshold for sole localization to the nucleus). Following the NLS, there is a run of 87 amino acids that is nearly perfectly duplicated once in the Type I GT1 and type III VEG strains, and twice in the Type II ME49 strain (ToxoDB.org). Following the large repeated domain there is an approximately 24 amino-acid sequence that is imperfectly repeated five times, before a predicted unstructured C-terminus (based on Protein Disorder Prediction System; http://prdos.hgc.jp/cgi-bin/prediction.cgi). These repeated sequences are consistent with conserved domains observed in other effector proteins, such as GRA16 and GRA24, where they have been reported to play a role in binding their cognate interacting partners. Additionally, the unstructured, serine-rich C-terminal region is consistent with a hypothesis that unstructured regions act as dynamic regions able to accommodate more signaling partners, as well as allowing for unfolding in order to cross the PVM (5). Thus the DNA sequence of *TGGT1_239010* suggests it encodes a protein that is exported, does not get stuck in a membrane, and would localize to the host nucleus.

**Figure 1.**
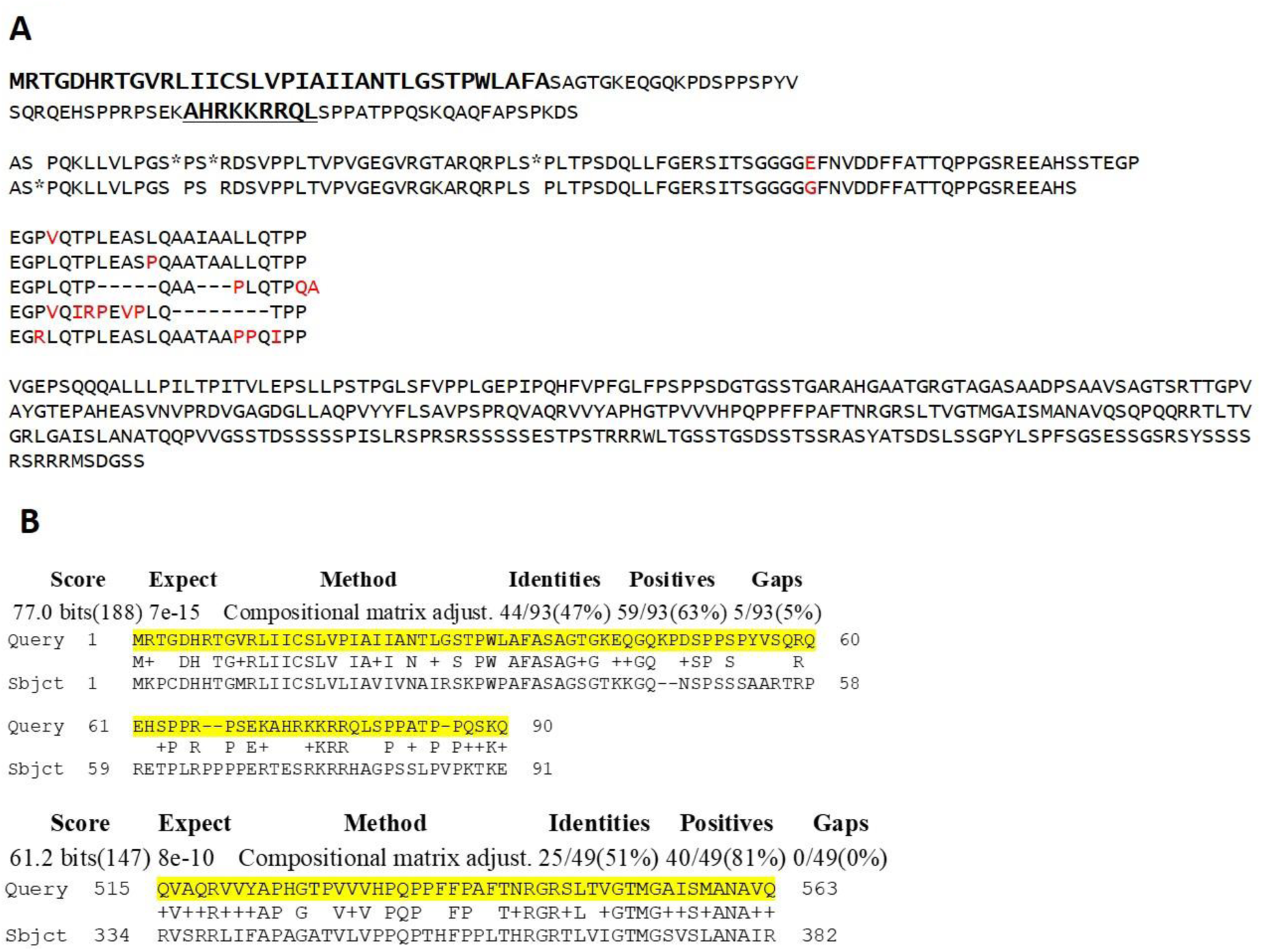
Predicted amino acid sequence and homology of TGGT1_239010. **A**. *TGGT1_239010* codes for a 685 amino acid protein that contains a predicted signal peptide (bold lettering) but no transmembrane domains internal to the protein, consistent with predictions for other effectors originating in dense granules. There is also a predicted nuclear localization signal (underlined bold) and several repeated domains of unknown function but repetitive structure. Red letters indicate discordance between the repeats and asterisks (*) indicate serine residues previously identified as being phosphorylated after secretion. Spaces have been added to make the repeated domains more clear. **B**. TGGT1_239010 is highly dissimilar to its orthologue BN1204_015825 in *Neospora caninum*. NCBI Blast was used to compare TGGT1_239010 (Query) to the *Neospora* proteome, and the only similarities found were within the two displayed regions of BN1204_015825 (Sbjct), including the predicted signal peptide and a segment of ~50 amino acids toward the C-terminus. Numbering indicates the amino acid position relative to the N-terminus in each predicted protein.

NCBI BlastP identified an orthologue, BN1204_015825, in *Neospora caninum* but with only two short regions of significant similarity: 47% identity (63% similarity) over the first 90 amino acids, mostly at the extreme N terminal region corresponding to the predicted signal peptide, and 51% identity (81% similarity) over a stretch of just 49 amino acids in the disordered, C-terminal region (Fig. 1B). This argues that TGGT1_239010 is under strong positive selection and that the function of the *Neospora* orthologue is likely very different from TGGT1_239010.

To begin characterization of *TGGT1_239010*, we first cloned its open reading frame, 5’-untranslated region and, to avoid overexpression artifacts, its predicted promoter into the plasmid pGRA and appended a sequence encoding a C-terminal hemagglutinin (HA) tag. Extracellular tachyzoites expressing this construct, referred to as 239HA, were stained with antibodies for HA and for GRA7, as a well-characterized marker for dense granules. Colocalization in discrete puncta was generally but not universally observed (Fig. 2A), consistent with TGGT1_239010 being a dense granule protein or a member of the “dense granule-like” proteins that have been observed for GRA16/MYR1/TgIST (2, 9, 11).

**Figure 2.**
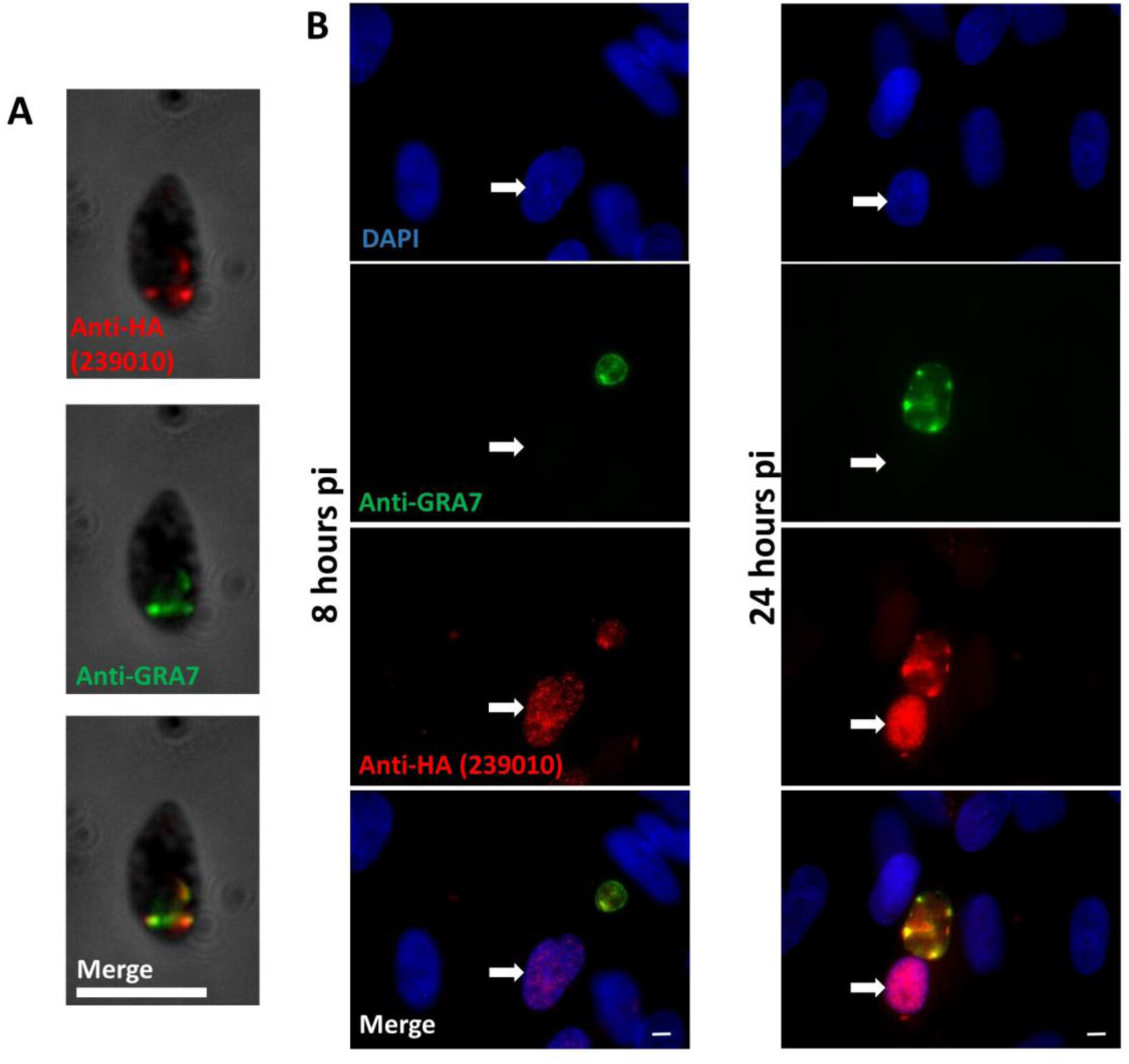
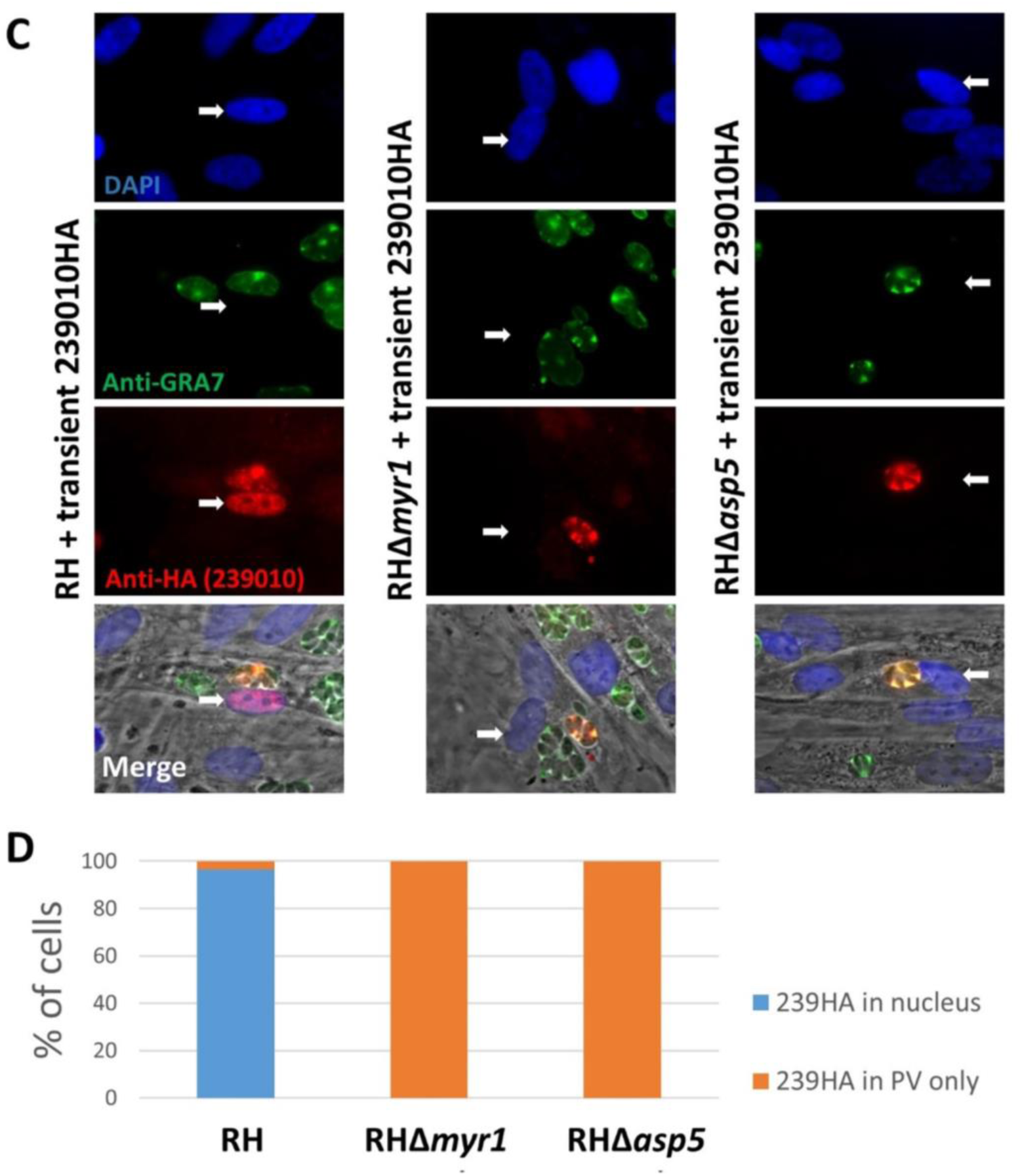
TGGT1_239010 is a dense granule protein that traffics to the nucleus of infected cells in a MYR1-and ASP5-dependent manner. **A.** Immunofluorescence assay (IFA) using anti-HA antibodies (red) to stain extracellular RH tachyzoites expressing TGGT1_239010 with an HA tag controlled by its endogenous promoter showing colocalization with known dense granule protein GRA7 (green) at discrete puncta within the parasite. White scale bar is 5 μm. **B.** IFA as for part A but showing RH tachyzoites infecting human foreskin fibroblasts (HFFs) 8 or 24 hours post-infection (pi) and DAPI staining to show the host nuclei. The white arrow points to the host nucleus in the infected cell. **C.** IFA as for part B but showing representative HFFs infected for 20 hours with wild type (RH), RHΔ*myr1*, or RHΔ*asp5* parasites transiently transfected with a construct expressing HA-tagged TGGT1_239010. TGGT1_239010 traffics to the host nucleus in RH-but not RHΔ*myr1*-or RHΔ*asp5*-infected HFFs. **D.** Quantification of the IFA data from the experiments shown in panel C. “% of Cells” is the percentage of infected host cells showing nuclear or PV staining with anti-HA antibody after infection with the indicated parasite line, n=28 (RH), n=41 (RHΔ*myr1)*, n=29 (RHΔ*asp5)*. Data shown are representative of two independent experiments that gave similar results.

When the same tachyzoites were used to infect human foreskin fibroblasts (HFFs) for 8 hours, 239HA localized within the parasitophorous vacuole, again with strong overlap with GRA7 (Fig. 2B). At 24 hours post-infection (hpi), it was apparent from the colocalization with GRA7 that the 239HA within the PV is largely located in the spaces between the parasites. More importantly, at both 8 and 24 hpi, 239HA also localized to the infected cell’s host nucleus, demonstrating that it is exported out of the vacuole and suggesting that the predicted NLS is functional. Comparing the results for 8 and 24 hpi, staining of 239HA increased over time in both the parasitophorous vacuole and nucleus, as previously noted for dense granule proteins (9).

All dense granule proteins known to localize to the host cell nucleus require the MYR machinery as well as an active ASP5 for translocation across the PVM, regardless of whether they are themselves cleaved by ASP5. To test if this was also the case for TGGT1_239010, we transiently transfected wild type RH, RHΔ*myr1*, and RHΔ*asp5* with a plasmid expressing 239HA. Expression of the transgene was confirmed in all three strains based on anti-HA staining in the parasitophorous vacuole, colocalizing with GRA7 (Fig. 2C); however, only in cells infected with the wild type RH was there observable 239HA in the host cell nucleus. Quantification of the results showed that in 96% of cells infected with the wild type parasites there was clear 239HA in the host nucleus, whereas no such nuclear staining was observed in the cells infected with the RHΔ*myr1* or RHΔ*asp5* mutants (Fig. 2D). Thus, as in the case of GRA24 (7), even though TGGT1_239010 has no TEXEL motif (Arg-Arg-Leu) and was not detected in an effort to identify all proteins cleaved by ASP5 (17), its export requires both this protease and intact MYR machinery.

### TGGT1_239010 Is Necessary and Sufficient for Host Cyclin E Induction

To determine if TGGT1_239010 is the effector controlling cyclin E (CCNE1) expression, we next compared the host cell response during infection with wild type parasites to the response during infection with parasites lacking TGGT1_239010. To create such a strain, we used CRISPR to target the single exon genomic sequence of *TGGT1_239010* for disruption by insertion of the *HXGPRT* gene (Supplemental Fig. 1A). Following selection for insertion of *HXGPRT*, we isolated clones and used PCR to amplify the *TGGT1_239010* locus using primers flanking the targeted insertion site. The results (Supplemental Fig. 1B) show the predicted change in size of the PCR product indicating disruption in the clones, and subsequent sequencing confirmed the disruptive insertion in *TGGT1_239010*.

With a confirmed knock-out in hand, we next performed RNASeq on mock-infected HFFs and HFFs infected for 6 hours with RH, RHΔ*239010* and RHΔ*myr1* at a high MOI of 5. We chose this MOI to be sure the vast majority of the HFFS are infected, and this 6-hour time point to allow parasites sufficient time to invade, establish a parasitophorous vacuole, and export TGGT1_239010 thereby altering host functions. We reasoned this would minimize the amount of downstream effects, with the goal of catching mostly the primary host targets of TGGT1_239010, should it prove to be the effector we sought. The results (Fig. 3A) showed that deletion of *TGGT1_23901*0 has a major impact on *Toxoplasma’s* ability to modulate host cell transcription. Using a threshold of at least 2.5-fold difference in a host gene’s transcript levels, we observed that 602 annotated genes were upregulated in cells infected with wild type RH that were not upregulated in cells infected with RHΔ*239010* (Supplemental Table 1). The relative expression level of these genes is also shown for HFF infected with RH*Δmyr1* and those that were mock-infected.

**Figure 3.**
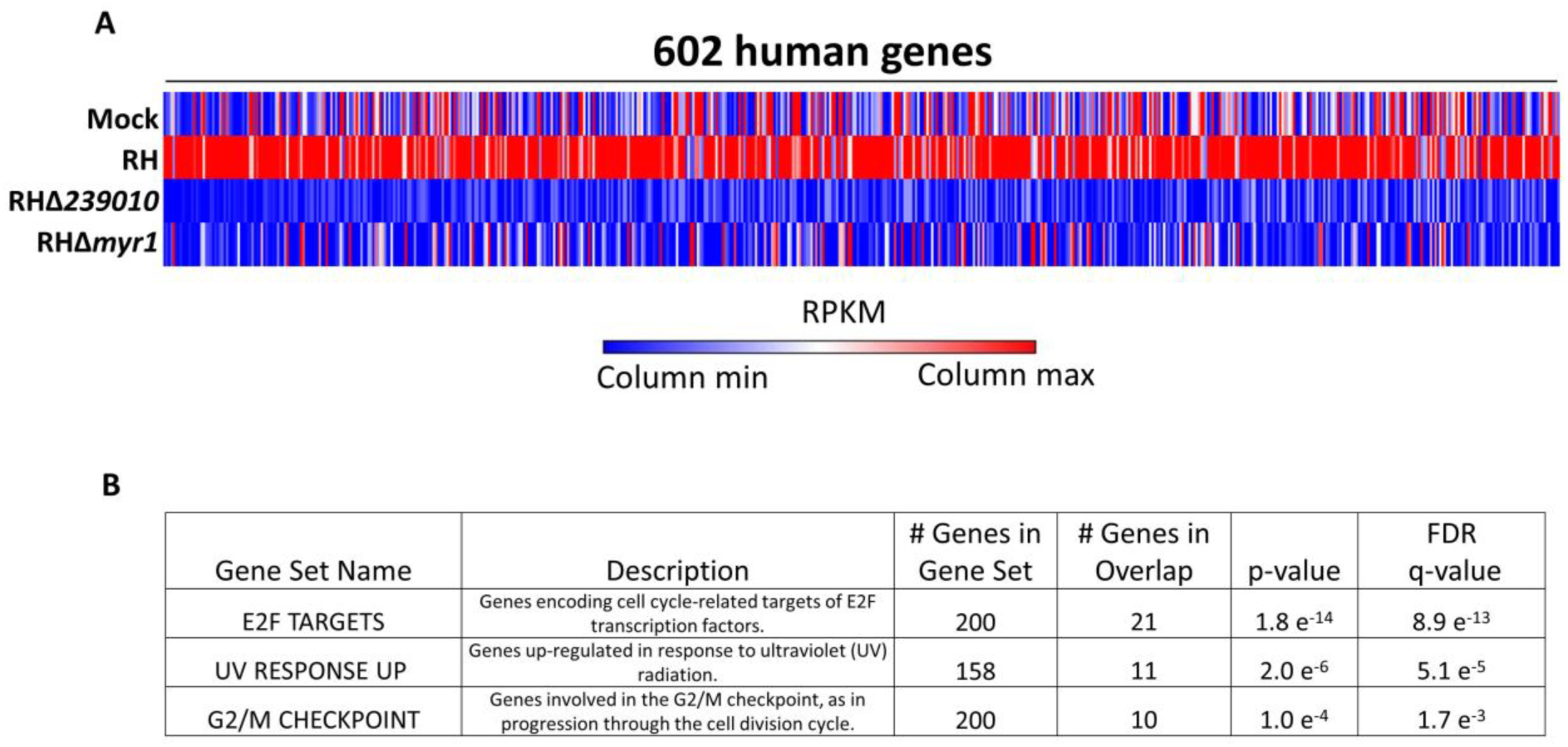
Disruption of Toxoplasma *TGGT1_239010* results in a parasite unable to alter the expression of genes associated with host cell cycle. **A**. Graphical representation of RNASeq of HFFs mock-infected or infected with the indicated *Toxoplasma* strains 6 hours post-infection. Values shown are the average relative RPKM from two independent experiments with red and blue being the highest and lowest values, respectively, for each gene. Shown are the 602 human genes that were annotated and showed a minimum 2.5-fold up-regulation in cells infected with RH-wild type vs. RHΔ*239010*. The values for these same genes in mock-infected HFF and in cells infected with RHΔ*myr1* are shown for comparison. Gene names and RPKM values can be found in Supplemental Table 1. **B.** Gene set enrichment analysis for the collection of host genes that are differentially up-regulated (p-value<0.0001) in HFFs infected with RHΔ*239010* vs. RH-wild type. Specific genes in each gene set can be found in Supplemental Table 2.

Gene Set Enrichment Analysis (GSEA) was then applied to the set of genes that were differentially expressed between RH-infected and RHΔ*239010-*infected HFFs, using a p value threshold of 1 x 10^−4^ to identify any gene sets that were significantly affected. The results (Fig. 3B) showed that the affected genes were most significantly enriched for genes impacted by the E2F family of transcription factors, having a p value of 1.8 x 10^−14^. We also observed significant effects on “ultraviolet radiation response” as well as “G2/M checkpoint” gene sets, albeit at lower p values of 2.0 x 10^−6^ and 1.0 x 10^−4^, respectively. E2F transcription factors are generally known to initiate the G1/S transition by an upregulation of cyclin E amongst other proteins, while UV response and G2/M checkpoint genes are involved in cell cycle checkpoints associated with both DNA replication and proceeding to mitosis. In total, GSEA analysis (http://software.broadinstitute.org/gsea/index.jsp) identified 21 genes that were targeted by E2F molecules, many of which are well known as parts of the pre-replication complex initiating DNA synthesis: MCM2, 3, 4, 5, and 7 as well as the DNA polymerase POLE (Supplemental Table 2).

Given the limited number of gene sets that appeared to be affected by the deletion of *TGGT1_239010*, as well as the fact that it ultimately is found in the host cell nucleus, it seemed unlikely that this protein was part of the effector translocation machinery, rather than being an effector itself. Nevertheless, and to confirm this, we tested whether the export of a different, known effector molecule, GRA16, was impacted by the loss of the *TGGT1_239010* gene. The results (Supplemental Fig. S2) showed that GRA16 accumulates normally in the nucleus of cells infected with RHΔ*239010* when transiently transfected with an HA-tagged GRA16. Thus, TGGT1_239010 is not a part of the general translocation machinery.

To gain further information about the role of TGGT1_239010, we next ranked host genes by their fold-difference in expression between RH-WT-infected and RHΔ*239010*-infected HFFs and examined those that had at least 10 reads per gene in the RH-infected sample, a number large enough to be confident in differential expression (Fig. 4A). Because E2F transcription factors drive cyclin E expression, it was not surprising to find that both cyclin E1 (CCNE1) and E2 (CCNE2) were among the top 25 most-affected genes. We chose to focus on these cyclins because they are canonical downstream targets of the E2F transcription factors and they play crucial roles in cell cycle control. To validate and extend the RNASeq results, we examined the protein levels of cyclin E1 in host cells infected with wild type and RHΔ*239010* parasites. Consistent with the elevated expression of its mRNA levels, IFA revealed that infected HFFs show an upregulation of cyclin E1 expression dependent on the parasite being wild type for both MYR1 and ASP5, as well as having an intact *TGGT1_239010* gene (Fig. 4B). This observation was confirmed by Western blots of infected cell lysates probed with antibodies to cyclin E1; cyclin E1 was strongly upregulated in HFFs infected with wild type tachyzoites but not with tachyzoites lacking MYR1, ASP5, or TGGT1_239010 (Fig. 4C). To ensure that the disruption of the *TGGT1_239010* locus was responsible for the phenotype, we generated a complemented strain (RHΔ*239010::239010HA*) and tested its ability to rescue the phenotype. The results (Fig. 4D) showed that, indeed, the complemented strain was fully capable of inducing the upregulation of cyclin E1 to levels indistinguishable from those seen in cells infected with wild type RH. Based on this, we dubbed the *TGGT1_239010* locus *HCE1* for its impact on “host cyclin E” expression.

**Figure 4.**
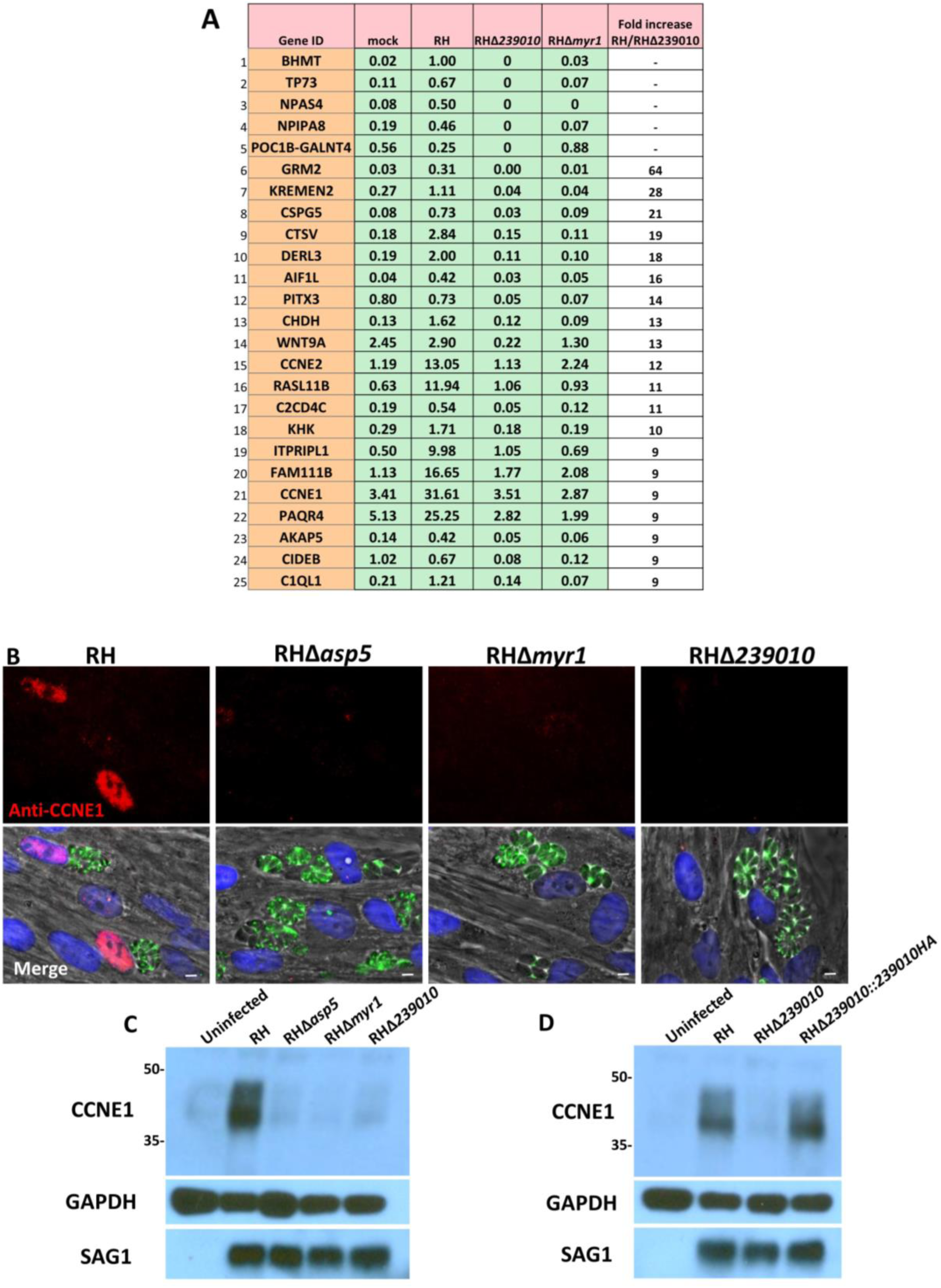
Cyclin E1 (*CCNE1*) and cyclin E2 (*CCNE2*) are among the top annotated genes whose up-regulation is dependent on TGGT1_239010. **A.** RPKM values for the top 25 human genes that have a minimum of 10 reads in both RH-infected samples and whose up-regulation is most dependent on TGGT1_239010. Genes are sorted based on the ratio of their expression during wild type infection to that during infection with RHΔ*239010*. RPKM values for cells infected with RHΔ*myr1* are shown for comparison. For instances where there was no expression of the gene in RHΔ*239010*-infected cells, genes with highest expression in RH-infected cells are listed first. **B.** IFA was used to assess the expression of host cyclin E1 (red) in the nuclei (blue) of cells infected with the indicated strains (green) for 20 hours. White scale bar is 5 μm. **C**. Western blot of cyclin E1 expression from HFFs infected for 20 hours with the indicated strains of *Toxoplasma*. Results from the same blots probed with antibodies for host GAPDH and parasite SAG1 are shown as loading controls. Size markers (kDa) are shown to the left. Blot shown is representative of three independent replicates. **D.** As for the Western blots in panel C but with the addition of a lysate from HFFs infected with a RHΔ*239010* strain carrying an ectopically located copy of the wild type *TGGT1_239010* gene including a sequence encoding a C-terminal HA-tag (RHΔ*239010::239010-*HA). Blot shown is representative of three independent replicates.

### HCE1 is Necessary for Efficient Parasite Growth in HFFs

To assess the importance of HCE1’s function in parasite growth, we measured plaque area using wild type and Δ*hce1* mutant parasites. The results (Fig. 5) show that plaques of tachyzoites lacking *HCE1* were ~65% the area of wild type plaques at day 7, and complementation with a copy of *HCE1HA* returned these plaques to their wild type size. Thus, HCE1 confers a growth advantage *in vitro*, even in HFFs which lack many immune defenses, consistent with an impact on host cell cycle control rather than interfering with known immune responses.

**Figure 5.**
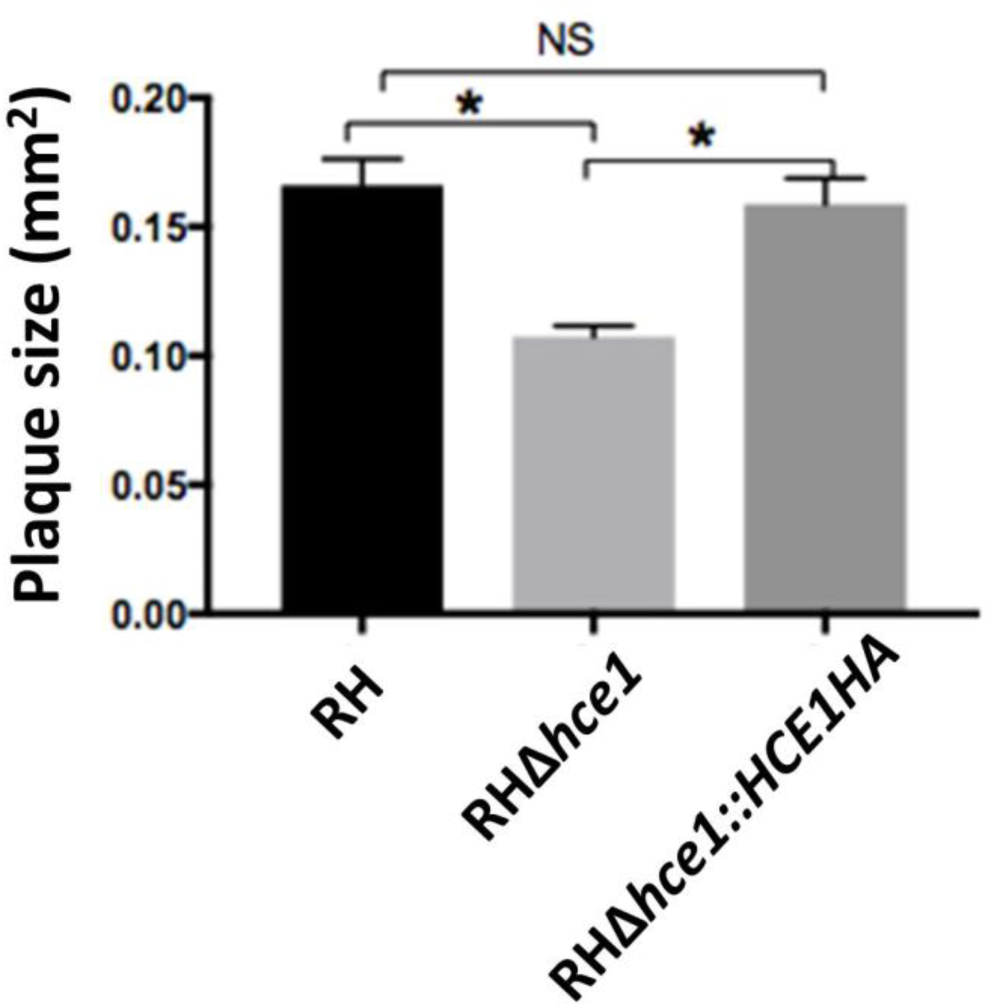
Growth of parasites in HFFs is retarded by the loss of HCE1. The indicated strains were used to infect HFFs and the plaque size was measured after 7 days of growth. Average plaque size was assessed by ImageJ for n=31 (RH), n=38 (RHΔ*hce1*), n=30 (RHΔ*hce1::HCE1HA*). Data are representative of 3 independent biologic replicates and error bars indicate the Standard Error of the Mean. Significance was determined by two tailed t test with * p<0.0001; NS, not significant (p>0.05).

### HCE1 Binds Host E2F/DP1 Heterodimers that Control Host Cell Cycle

To shed light on how HCE1 functions, we next sought to determine its binding partners. To do this, we used anti-HA antibodies to immunoprecipitate HA-tagged HCE1 from cells infected for 24 hours with RHΔ*hce1::HCE1HA* tachyzoites under conditions where associating proteins are likely to remain intact (IP #1). As a control for proteins that might be precipitated with the anti-HA beads nonspecifically, we also performed this experiment with HFFs infected with an untagged RH strain. We performed this same experiment a second time under more stringent conditions, disrupting weak interactions and releasing proteins from membranes with an additional sonication step and comparing the results to an untagged RH strain (IP #2). Mass spectrometry (MS) of the proteins enriched by the anti-HA immunoprecipitation in both experiments was performed and the results were ranked by the ratio of the relative number of spectral counts found for a given protein in the HCE1HA lysates compared to RH, with a nominal one count added to all values to facilitate the mathematical calculation.

When we searched the IP-MS data for *Toxoplasma* proteins associating with HCE1, we observed eleven to be highly enriched (an enrichment score ≥5 in both experiments; Supplementary Table 3). Among these was MYR1, that we know is necessary for HCE1 translocation, possibly explaining its association, and other GRA proteins that we know are secreted into the PV (GRA28 and GRA44) of which at least one (GRA28) is also translocated into the host cell (Supplemental Table 3) (17, 18). One or more of these proteins may be additional players in the trafficking of HCE1 across the PVM and into the host nucleus but we have not pursued this further because, as described further below, we find HCE1 can perform its major functions independent of all other *Toxoplasma* proteins.

When the immunoprecipitation data were examined for human proteins that appear to specifically associate with HCE1, we observed seventeen with enrichment scores greater than 2 in both experiments (Figure 6A; a complete list is in Supplemental Table 3). The three proteins that were most strongly enriched were DP1, E2F3 and E2F4 which were identified with at least 7 spectral counts (and at least 6 unique peptides) in both immunoprecipitations with HCE1HA and 0 spectral counts in both immunoprecipitations with the negative control, RH. Note that E2F3 is expressed as two variants differing in the initiating methionine; E2F3a has an N-terminal extension of approximately 120 amino acids over E2Fb, and when we mapped unique peptide fragments to these two isoforms we found 12 out of 13 fragments map to the common region but 1 peptide fragment, with 1 spectral count in each IP, maps to the region unique to E2F3a (Supplemental Figure 3). Thus, while the data are unambiguous in indicating the presence of E2F3, we cannot conclude whether E2F3b is present or absent but would argue that the E2F3a variant, at least, is present.

**Figure 6.**
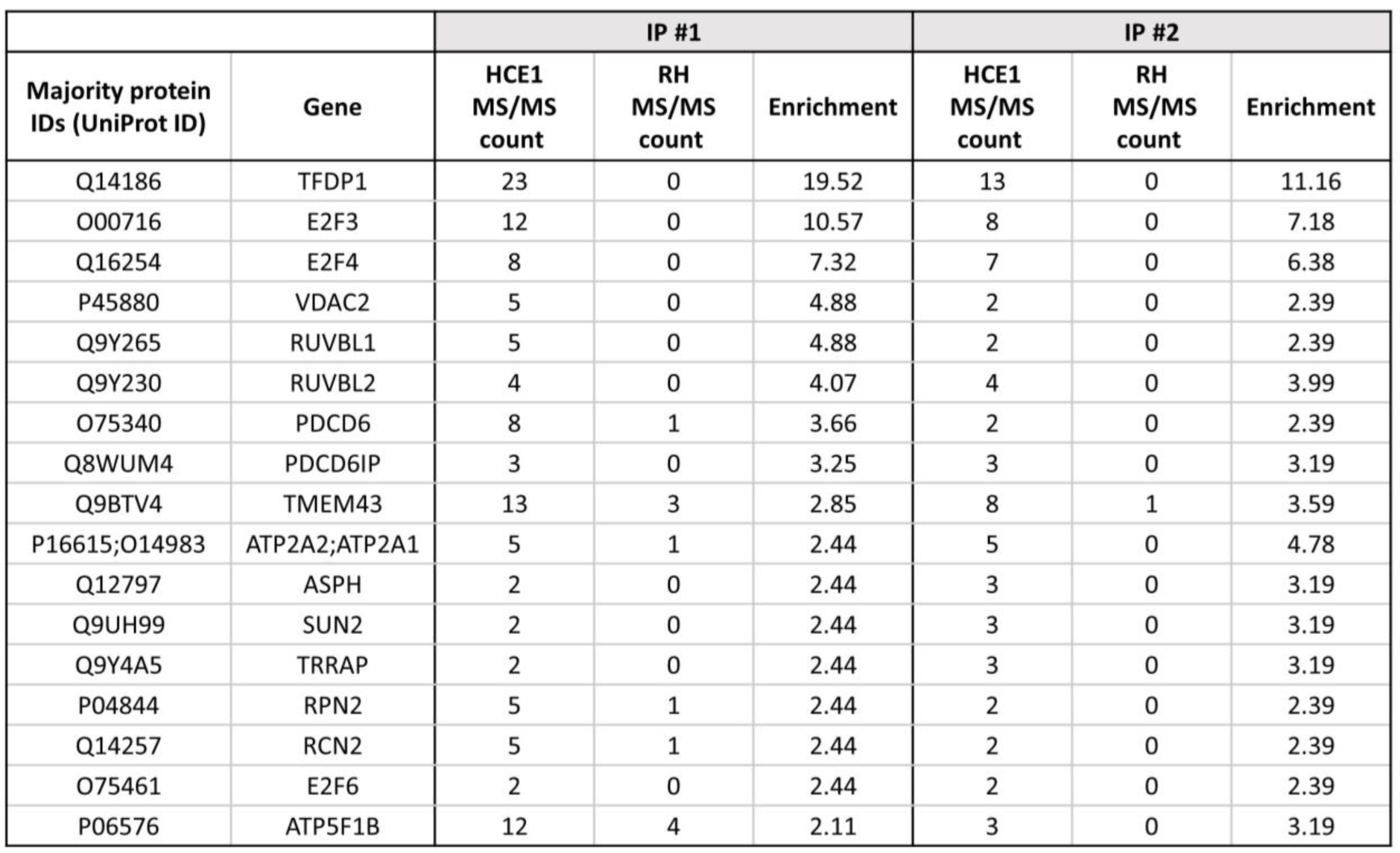
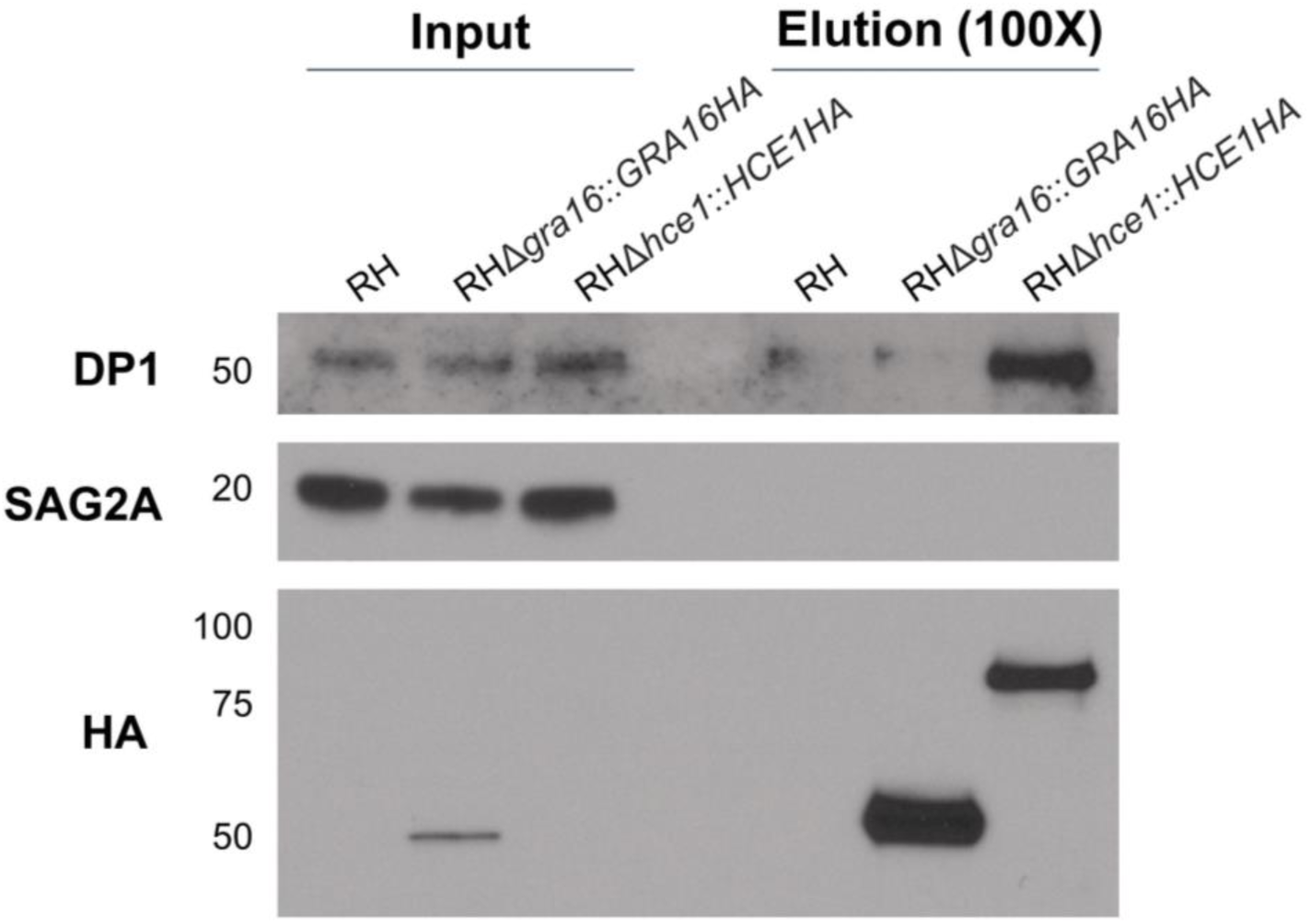
HCE1 specifically associates with host TFDP1 and E2F. **A**. Anti-HA magnetic beads were used to immunoprecipitate proteins from HFFs infected for 24 hours with RH wild-type or RHΔ*hce1::HCE1HA*. Mass spectrometry was performed on the resulting material and the number of spectral counts was determined for all detectable proteins (IP #1). The experiment was repeated with the same conditions except the lysate was sonicated prior to immunoprecipitation to ensure release of all proteins that might be trapped in membranous material (IP #2). The results for both experiments were ranked according to the enrichment of spectral counts in the HCE1-HA-expressing strain relative to the wild-type RH control after adding a nominal single count to all results, thereby enabling a ratio to be determined, and accounting for the total spectral counts identified in each experiment (Enrichment). Displayed are the majority protein IDs, i.e the proteins which contain at least half of the peptides belonging to a protein group (grouping of proteins which cannot be unambiguously identified by unique peptides), the corresponding gene for those proteins, and the corresponding number of spectral counts (MS/MS count) for all human proteins with an enrichment score greater than 2 in both of the experiments. The top three enriched proteins in either experiment were DP1, E2F3, and E2F4. **B**. Lysates from a repeat of IP #2, including an additional control of parasites expressing an unrelated, HA-tagged effector, GRA16HA (RH*Δgra16::GRA*16HA), were resolved by SDS-PAGE, blotted and probed with antibodies to DP1, SAG2A (as a specificity and parasite input control) and anti-HA (to show efficient immunoprecipitation of the relevant HA-tagged protein). Approximately 100-fold more starting material is represented in the eluate than the input, and hence the stronger bands for the HA-tagged material in the eluate relative to input. DP1 was specifically enriched in the immunoprecipitation from the HCE1HA-expressing parasites. Blot indicates a single replicate.

Other components of the DP1-E2F transcription complex were also observed in the HCE1HA-associating material: E2F6, TRRAP (a histone acetyl transferase that associates with DP1-E2F), and RUVB1/RUVB2 (members of the AAA+ family of ATPases associated with various cellular activities and reported binders of E2F1 (19, 20). Six of the remaining ten proteins in the list had one or more spectral counts in the negative control making their specific association with HCE1 questionable. To validate the association of HCE1 with the E2F/DP1 transcription complex, we focused on DP1, which is the stably associated “dimerization partner” (hence DP1) of almost all E2Fs. For this validation, we repeated the infection and immunoprecipitations and included parasites expressing GRA16HA as an additional specificity control, since this protein is also present in the infected host cell nucleus (9). We then probed the eluates of the immunoprecipitation with an anti-DP1 antibody, and antibodies to SAG2A (as a loading control for input material) and HA (to verify the immunoprecipitation was successful; Fig. 6B). Although expression of HCE1HA was not detected in the input material, following immunoprecipitation and enrichment with anti-HA, HCE1HA was observed to migrate at ~80 kDa, consistent with its predicted size of 68.4 kDa (with HA tag and after removal of signal peptide) and allowing for the fact that it is phosphorylated at a minimum of 4 positions (Fig. 1A, (16)). Importantly, the results showed that DP1 is substantially enriched in the material coprecipitating with HCE1HA relative to the wild type and GRA16HA-expressing negative controls, confirming the specific association of DP1 with HCE1HA. Although we cannot discern from these results whether this association is direct or indirect (e.g., HCE1HA binds to E2F3 and E2F4 and, thereby, DP1), they do provide the beginning of a mechanistic explanation for how HCE1 regulates cyclin E, as discussed further below.

### HCE1 is Sufficient to Upregulate Cyclin E during Infection With *Neospora caninum* and When Expressed in Uninfected HFFs

Given the evidence from previous transcriptomic experiments with bovine trophoblast cells (14), plus the extreme divergence reported by NCBI BlastP between HCE1 and the *Neospora* orthologue BN1204_015825, we hypothesized that *Neospora caninum* would not induce cyclin E1 in human cells. This was tested and confirmed to be the case (Fig. 7A). To ask if HCE1 would be sufficient to confer the ability to upregulate cyclin E1 on its Apicomplexan cousin, we first generated a HCE1-expressing strain of *Neospora* and tested if the introduced protein would be exported to the host nucleus. As shown in Figure 7B, the HCE1 was indeed translocated to the host nucleus and so we then asked if the resulting strain would induce cyclin E. The results (Fig. 7C) showed that HFFs infected with this engineered strain show strong upregulation of cyclin E. Western blots (Fig. 7D) confirmed this finding on a population level. Collectively, these results indicate that *Neospora* has the ability to produce, process and export a fully functional HCE1 and that HCE1’s activity is not dependent on other effectors specific to *Toxoplasma*.

**Figure 7.**
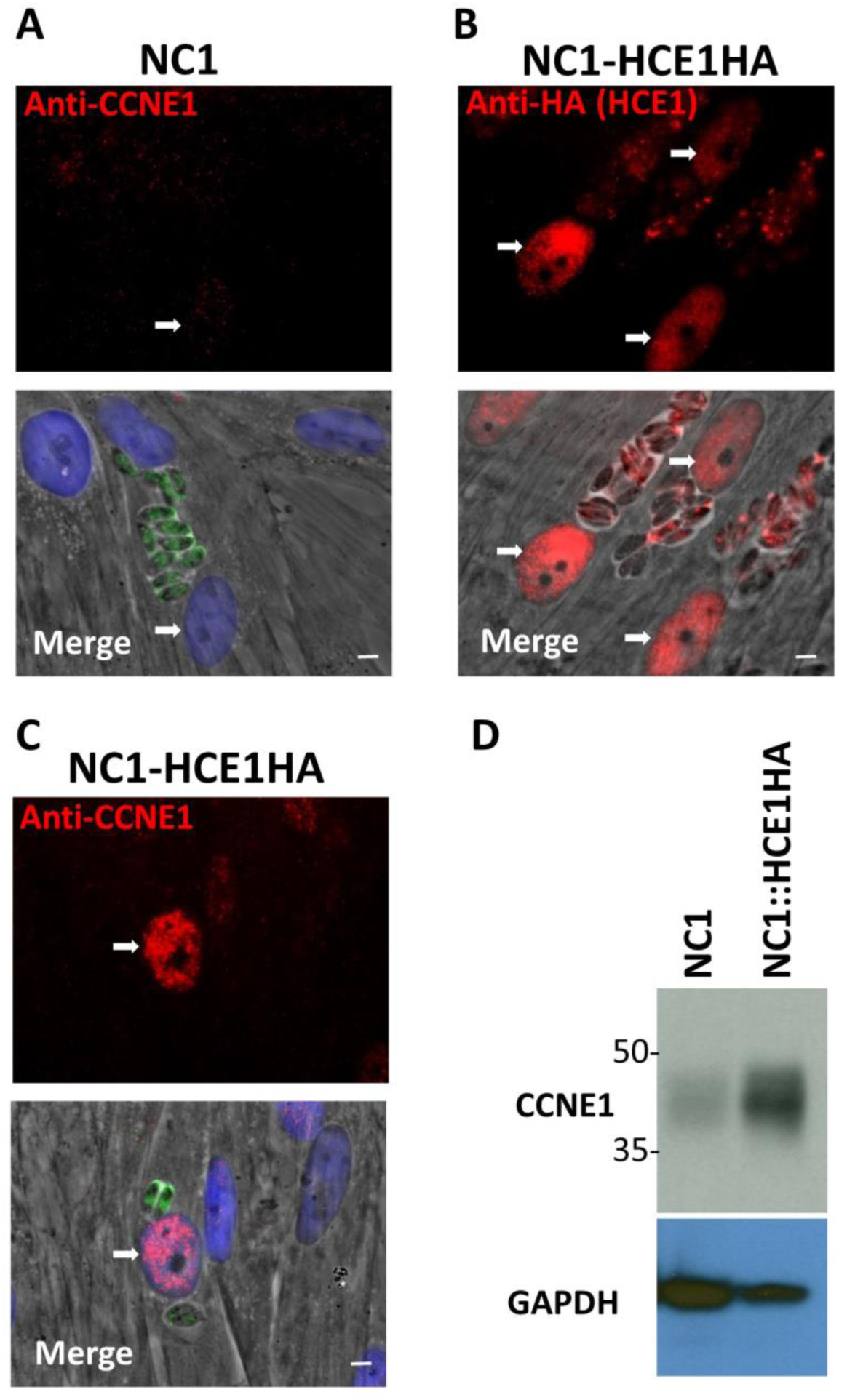
Expression of HCE1 in *Neospora* allows it to control host cyclin E1. **A.** IFA with anti-CCNE1 of HFFs infected with wild-type *Neospora caninum* NC1 showing no CCNE1 induction in the infected host cell’s nucleus (white arrows). White scale bar is 5 μm. **B.** As for Panel A except the IFA is with anti-HA of HFFs infected with NC1 stably transfected with *pGRA-TGGT1_239010HA* showing the transfected NC1 express, export, and localize HCE1 to the infected cell’s nucleus. **C.** As for Panel B except that the IFA is with anti-CCNE1 showing efficient CCNE1 induction in the infected host cell nuclei. **D**. Lysates of HFFs infected with the indicated NC1 strain for 20 hours were resolved by SDS-PAGE, blotted and probed with antibodies to CCNE1 or GAPDH (as a loading control). Blot is representative of three independent experiments.

We next tested whether HCE1 is sufficient to induce cyclin E on its own, with no infection and no other parasite proteins present. To do this, we cloned a truncated version of the HCE1 open reading frame, lacking the predicted signal sequence for export, into the human expression vector pcDNA under the CMV promoter and used Lipofectamine LTX to transfect this into subconfluent HFF. Subconfluent HFF were utilized in this experiment because the Lipofectamine LTX yields a higher transfection efficiency than confluent HFF. Neither the transfection procedure alone (No DNA) nor expression of an irrelevant gene (GFP) under the CMV promoter induced robust cyclin E1 (Fig. 8A), although a small fraction (~2.5%) of cells did express detectable cyclin E1 as expected for these subconfluent HFF cultures containing replicating cells (Fig. 8B). Transfection of the plasmid pcDNA-HCE1HA, however, resulted in strong expression of HCE1, a majority of which was concentrated within the cell nucleus (Fig. 8A). Importantly, >90% of cells expressing the HCE1 transgene exhibited this robust cyclin E1 induction (Fig. 8B).

**Figure 8.**
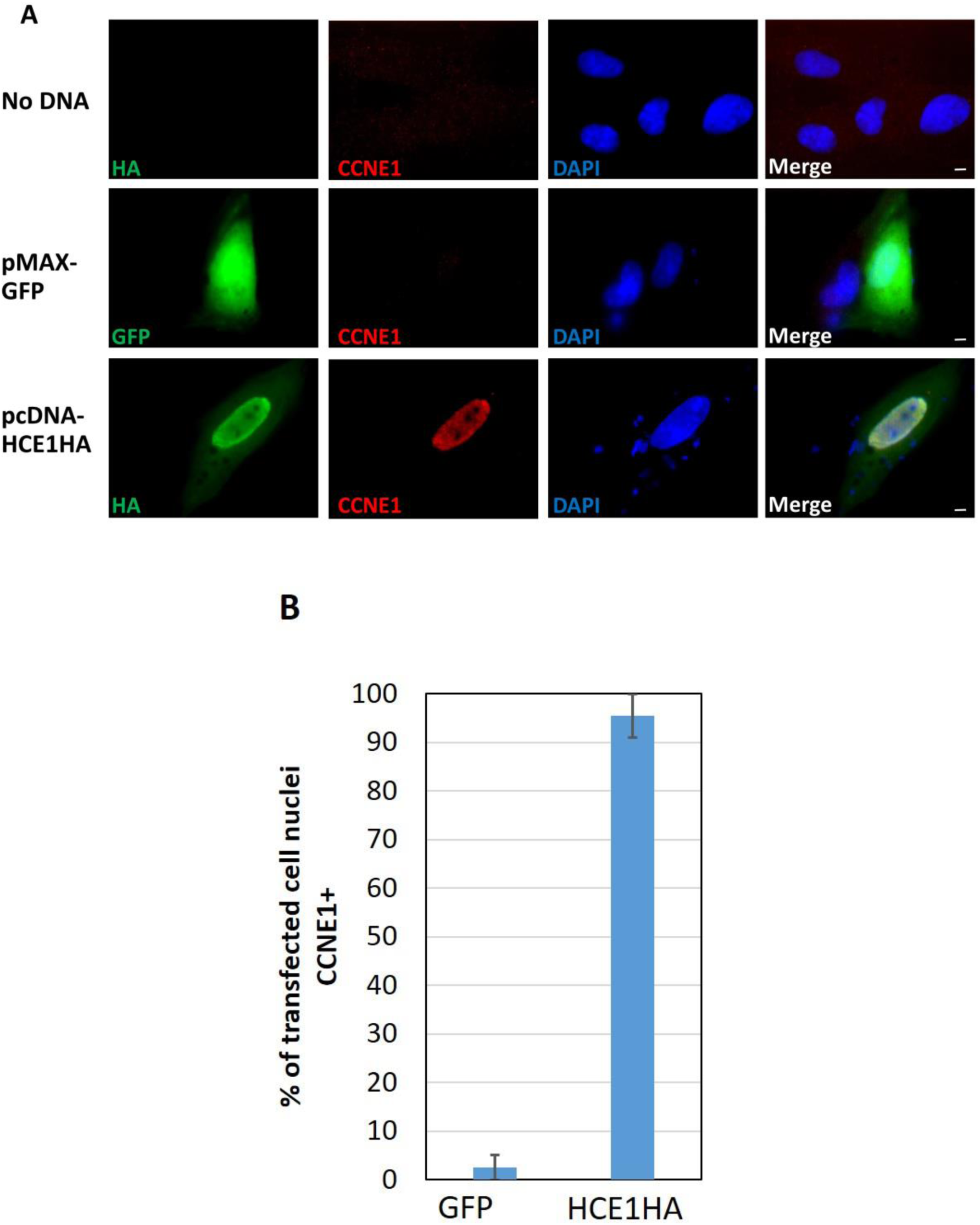
Expression of HCE1 in HFFs, without infection, is sufficient to up-regulate cyclin E1 at 20 hrs. **A.** HFFs were mock-transfected with no DNA or transfected with the human expression plasmid pMAX-GFP or pcDNA-HCE1HA, as indicated, and then assessed by IFA for CCNE1 expression at 20 hours. White scale bar is 5 μm. **B.** Quantification of HFF cultures shown in Panel A showing the mean percentage of host nuclei in cells expressing the indicated transgene that were positive for CCNE1. Data shown are averages of the assessment of two independent experiments and error bars indicate Standard Error of the mean.

## Discussion

We show here that HCE1 is an effector protein that is both necessary and sufficient for robust upregulation of cyclin E in the host, independent of all other parasite proteins. Mechanistically, once HCE1 arrives in the host cell nucleus it appears to function via a direct interaction with E2F/DP1 heterodimers, most prominently in the confluent HFFs studied here, those comprised of E2F3 and E2F4. We also detected binding to E2F6, the E2F-associating histone acetyl transferase TRRAP (20) and two AAA+ proteins that participate in chromatin remodeling, RUVB1 and RUVB2 (19). When DP1 is bound to E2F3, the complex is considered nominally “activating”, while when bound to E2F4 or E2F6 the complex is generally considered “inhibitory” (21, 22); however, this characterization depends heavily on other members of the transcription complex. For example, when cells are in the resting G_0_ state or the early G_1_ phase, before commitment to enter the cell cycle, the Retinoblastoma (RB) family of proteins (RB1, RBL1, RBL2) associate with E2F/DP1 dimers to maintain repression at the promoters of cyclin E and other E2F targets. Without RBL1 or RBL2, E2F4 has the potential to function as an activating transcription complex (23). Unlike what has been reported for adenovirus E1A, HPV-16 E7 proteins and polyoma viruses’ large T antigen, which can all bind RB to release active E2F (24-26), we did not observe binding of HCE1 with any of the RB family of proteins. Collectively, our results instead suggest a model in which HCE1 retains DP1/E2F3 and DP1/E2F4 in an active state, tied to the coactivator TRRAP, and without the suppressive effects of the RB pocket proteins.

Several publications have described the active interference of *Toxoplasma* tachyzoites with host cell cycle machinery (1, 2, 27, 28), including in MYR1-dependent ways (12). The results presented here for HCE1 likely provide at least a partial explanation for these effects since among the E2F-regulated genes whose expression is HCE1-dependent, we see several genes related to the pre-replication complex such as the minichromosome maintenance genes (MCM), DNA polymerase E and Origin Replication complex subunit 1 (ORC1). Previous observations that *Toxoplasma*-infected host cells are simultaneously induced from G1 into S and yet halted at the G2/M boundary have suggested that multiple effectors are likely in play; i.e., both an activator of the cell cycle and, later, an inhibitor. GRA16 is known to impact p53 (9) and GRA24 impacts p38 MAP Kinase signaling (29), both of which affect cell cycle. In the case of GRA16, a strong p53 signature combined with data demonstrating a decrease in cyclin B are consistent with it being one of the effectors to stall the cell cycle in G2 (9). This may provide a halt to the effect that is launched by HCE1, preventing a cell from actually dividing. Determining exactly how each effector’s function interfaces with the others in controlling cell cycle and what specific benefit this disruption provides to the growth of tachyzoites (evident from the fact that Δ*hce1* parasites produce significantly smaller plaques than wild type parasites), will require substantial further study. It may well be that launching a cell into an abortive process of growth and division provides a wealth of nutrients for the parasites to use for their own growth, and metabolomic studies of wild type-and Δ*hce1*-infected cells will help reveal whether such might be occurring. Regardless, the finding here that HCE1 is the protein responsible for cyclin E upregulation reveals its role as a critical lynchpin in how *Toxoplasma* tachyzoites so capably co-opt infected host cells for their own purposes.

## Materials and Methods

### Parasite strains, culture and infections

The following strains were used in this study: *Toxoplasma* RHΔ*hxgprt* (2), RHΔ*myr1* (2), RHΔ*asp5* (30), RHΔ*gra16* (6), RHΔ*gra16::GRA16HA (6)* and *Neospora caninum* NC-1 (31). *Toxoplasma* and *Neospora* tachyzoites were propagated in human foreskin fibroblasts (HFFs) cultured in complete Dulbecco’s Modified Eagle Medium (cDMEM) supplemented with 10% heat-inactivated fetal bovine serum (FBS; HyClone, Logan, UT), 2 mM L-glutamine, 100 U ml^−1^ penicillin and 100 µg ml^−1^ streptomycin at 37 °C with 5% CO_2_. These strains are available by contacting the authors.

The HFFs were obtained from the neonatal clinic at Stanford University following routine circumcisions that are performed at the request of the parents for cultural, health or other personal medical reasons (i.e., not in any way related to research). These foreskins, which would otherwise be discarded, are fully de-identified and therefore do not constitute “human subjects research”. Prior to infection, parasites were scraped and lysed using a 27 G needle, counted using a hemocytometer and added to HFFs at the stated multiplicity of infection. “Mock” infection was done by first syringe-lysing uninfected HFFs, processing this in the same manner as done for the infected cells, and then adding the same volume of the resulting material as used for infections.

### Transfections

All transfections were performed using the *BTX* EMC600 *Electroporation System* (Harvard Apparatus). Tachyzoites were mechanically released in PBS, pelleted, and resuspended in solution for transfection. After transfection, parasites were allowed to infect HFFs in DMEM. Transfections were performed using 5-10 x 10^6^ parasites and 3-10 µg DNA in Cytomix (10 mM KPO_4_, pH 7.6, 120 mM KCl, 5 mM MgCl_2_, 25 mM HEPES, 2 mM EDTA, 150 µM CaCl_2_).

### Immunofluorescence assay (IFA)

Infected cells grown on glass coverslips were fixed using 4% formaldehyde in PBS for 20 min. Samples were washed once with PBS and blocked using 3% BSA in PBS for 1 hour at room temperature (RT). Cells were permeabilized with 0.2% Triton-X100/3% BSA/PBS for 10 min at RT. Cyclin E was detected with mouse monoclonal antibody HE111 (Santa Cruz Biotechnology), GRA7 was detected with rabbit polyclonal anti-GRA7 serum, and HA was detected with rat monoclonal antibody 3F10 (Roche). Secondary anti-mouse, -rat and -rabbit antibodies were used conjugated to Alexafluor 488 and 594. Vectashield with DAPI stain (Vector Laboratories) was used to mount the coverslips on slides. Fluorescence was detected using wide-field epifluorescence microscopy and images were analyzed using ImageJ or FIJI. All images shown for any given condition/staining in any given comparison/dataset were obtained using identical parameters.

### Gene disruption

The RHΔ*HCE1* and RHΔ*gra16* strain were generated by disrupting the corresponding gene loci using CRISPR-Cas9 and selecting for integration of a vector encoding hypoxanthine-guanine phosphoribosyltransferase (*HXGPRT)* using drug selection. Specifically, the pSAG1:U6-Cas9:sgUPRT vector (32) was modified by Q5 site-directed mutagenesis (NEB) to specify a sgRNA targeting *TGGT1_239010*. The resulting sgRNA plasmid, dubbed pSAG1:U6-Cas9:sg239 was transfected into RHΔ*hpt* along with the previously described pTKO2 vector (33), which carries the *HXGPRT* gene flanked by *loxP* sites. Parasites were allowed to infect HFFs for 24 hrs, after which time the media was changed to complete DMEM supplemented with 50 µg/ml mycophenolic acid (MPA) and 50 µg/ml xanthine (XAN) for HXGPRT selection. The parasites were passed twice before being single cloned into 96-well plates by limiting dilution. Disruption of the gene-coding regions was confirmed by PCR using primers 239010F (GCACGAACCATAGAAAAGTAGGAA) and 239010R (AGTGGTCGCTGGCGTGCTGTT) and then sequencing of the amplification products.

### Complementation

The RHΔ*239010* strain (also referred to as RH*Δhce1*) was complemented ectopically with the pGRA-239010-HA (pGRA-HCE1HA) plasmid. Expression of *TGGT1_239010*/*HCE1* is driven off its natural promoter. To construct the pGRA-239010-HA plasmid, the *TGGT1_239010* promoter and open reading frame were amplified from RHΔ*hpt* genomic DNA using ACTAAAGCTTTTAGGCCAAAAACTGCACCCATCC and TAGTTTAATTAACTACGCGTAGTCCGGGACGTCGTACGGGTAGGAAGATCCGTCCGACATTCTTC primers. The resulting ~3.5 kb fragment was digested with HindIII and PacI restriction enzymes and cloned into the corresponding sites of the pGRA1 backbone. Five micrograms of the resulting vector, pGRA-HCE1-HA was co-transfected with 3 µg pSAG1:U6-Cas9:sgUPRT (32) into RHΔ*hce1* tachyzoites to create RHΔ*hce1::HCE1-HA*. Integration of the vector at the *UPRT* locus was enriched by selecting for resistance to 5 µM FUDR in DMEM after one lytic cycle. The resulting population was then cloned by limiting dilution and tested for HCE-HA expression by IFA.

### Gene expression in human foreskin fibroblasts

The *239010* open reading frame was cloned from genomic DNA isolated from RH-infected HFF. Primers TAGTGCGGCCGCATGTTTGCAAGCGCCGGAACGGG and TAGTCTCGAGCTACGCGTAGTCCGGGACGTCGTACGGGTAGGAAGATCCGTCCGACATTCTTC were used to amplify the reading frame after the signal sequence (beginning with the codons encoding PheAlaSerAla), add a C-terminal HA tag, and insert into pcDNA after digestion with NotI and XhoI. GFP was expressed from the pMAX-GFP plasmid. 1000-1500 ng of DNA was transfected into subconfluent HFFs using Lipofectamine LTX (Thermo) according to the manufacturer’s protocol.

### Western blotting

Infections for Western blot were performed at an MOI of 2:1. Parasites were syringe-lysed using a 27 G needle, counted and equivalent numbers of parasites were used for infections. Cell lysates were harvested 20 hours post-infection (hpi) and suspended in RIPA buffer with protease and phosphatase inhibitors (Roche, Thermo Fisher). Samples containing 10-20 µg protein were boiled for 20 min in sample buffer, separated by SDS-PAGE, and transferred to polyvinylidene difluoride (PVDF) membranes. Membranes were blocked in 5% milk in TBS supplemented with 0.2% Tween-20 for 1 hour at RT and then incubated 1 hr at RT with primary antibody in blocking buffer. Cyclin E1 was detected using mouse monoclonal antibody HE12 (Santa Cruz Biotechnology) and goat anti-mouse secondary antibody conjugated to horseradish peroxidase (HRP). GAPDH was detected with mouse monoclonal antibody 6C5 (Calbiochem). SAG1 levels were used to control for the levels of parasites within the infected cells and blots were stained with polyclonal rabbit anti-SAG1 followed by a secondary goat anti-Rabbit antibody conjugated to HRP. The HA epitope was detected using horseradish peroxidase (HRP)-conjugated HA antibody (Roche; catalog no. 12013819001 ab 3F10). SAG2 and DP1 were recognized by rabbit polyclonal anti-SAG2 (generated previously (34), and mouse monoclonal anti-DP1 (Santa Cruz Biotechnology, catalog no. sc-70990), respectively. For anti-DP1, a secondary antibody that recognizes the non-reduced form of mouse IgG was used (Abcam catalog no. ab131368). HRP was detected using enhanced chemiluminescence (ECL) kit (Pierce). Membranes were stripped between blots by incubation in stripping buffer (Thermo Fisher) for 10-15 min then washed 2 x 5 min with TBS-T.

### Plaque assay

HFFs were grown to confluency in a T25 flask, 200 parasites were added to each T25 and incubated for 7 days. The infected monolayers were washed with PBS, methanol fixed and stained with crystal violet. Plaque sizes were measured using ImageJ.

### RNA extraction, library preparation, and sequencing

HFFs were serum-starved for 24 hours before infection by growth in DMEM containing 0.5% serum. They were then infected with the indicated line of tachyzoites at an MOI of 5, and at 6 hpi, 1 ml TRIzol reagent (Invitrogen) was added to each T25 and the cells were scraped. Lysates were collected into RNAse/DNAse-free Eppendorf tubes and frozen at −20 °C. Total RNA was extracted following the manufacturer’s instructions, with some modifications. Briefly: frozen samples were thawed on ice and 0.2 ml chloroform was added to TRIzol suspensions, which were then mixed by inverting 10 times, and incubated for 5 min. Tubes were then spun at 12,000 rpm for 15 min at 4 °C. RNA in the aqueous phase was transferred into a fresh tube and 0.5 ml absolute isopropyl alcohol was added. Each tube was inverted three times and incubated at 4 °C for 10 min. They were then spun at 12,000 rpm for 20 min at 4 °C. After decanting the supernatants, RNA pellets were washed with 1 ml 75% ethanol. Tubes were mixed by inverting 10 times and then spun at 12,000 rpm for 20 min at 4 °C. Supernatants were removed and the RNA pellets were air-dried in open tubes for approximately 10 min. The RNA pellets were resuspended in 30 μl RNase-free DEPC-water. Multiplex sequencing libraries were generated with RNA Sample Prep Kit (Illumina) and samples were submitted to the Stanford University Functional Genomic Facility (SFGF) for purity analysis using the Agilent 2100 Bioanalyzer. Samples, having been barcoded to preserve identity, were then pooled for a single high-throughput sequencing run using the Illumina NextSeq platform (Illumina Nextseq 500). Infection and harvest were done twice, independently.

### RNASeq read mapping and differential expression analysis

Raw reads were uploaded onto the CLC Genomics Workbench 8.0 (Qiagen) platform for independent alignments against the human genomes (Ensembl.org/ hg19). All parameters were left at their default values. The number of total reads mapped to each genome was used to determine the RPKM (Reads Per Kilobase of transcript per Million mapped reads). Heat maps were generated using Gene E (https://software.broadinstitute.org/GENE-E/index.html).

### Gene Set Enrichment Analysis (GSEA)

GSEA, available through the Broad Institute at http://www.broadinstitute.org/gsea/index.jsp, was the enrichment analysis software used to determine whether defined sets of differentially expressed human genes in our experiment show statistically significant overlap with gene sets in the curated Molecular Signatures Databases (MsigDB) Hallmark gene set collection. We used the cutoff of p-value ≤10^−4^. The list of genes that are found in the gene sets presented are displayed in Supplemental Table 2.

### Data sets

The RNASeq data files have been deposited in GEO under accession number GSE122786. The mass spectrometry proteomics data have been deposited to the ProteomeXchange Consortium (http://proteomecentral.proteomexchange.org) via the PRIDE partner repository (35) with the dataset identifier PXD012103.

### Immunoprecipitations (IPs) for Mass Spectrometry Samples

IPs to identify HCE1-interacting proteins in HFFs were performed as follows. One 15-cm dish of HFFs for each infection condition was grown to confluence. HFFs were infected with either 15 x 10^6^ RH, RH?*hce1*::*HCE1HA*, or RH?*gra16*::*GRA16HA* parasites for 24 hours. Infected cells were washed 3 times in cold PBS and then scraped into 1 mL cold cell lysis buffer (50mM Tris (pH 8.0), 150mM NaCl, 0.1% (v/v) Nonidet P-40 Alternative [CAS no. 9016-45-9]) supplemented with complete protease inhibitor cocktail (cOmplete, EDTA-free [Roche]). Cell lysate was passed 3 times through a 25 G needle, followed by 3 times through a 27 G needle to break up the cells. For IP #2 only, the cell lysate was then subjected to sonication on ice (Branson Sonifier 250, with 3 pulses of 10 s at 50% duty cycle and output control 2). Cell lysates were spun at 1000 × *g* for 10 min to remove insoluble material and unlysed cells. Lysates were added to 100 µL magnetic beads conjugated to anti-HA antibodies (Pierce) and incubated overnight rotating at 4 °C. Unbound protein lysate was removed, and the anti-HA magnetic beads were then washed 10 times in cell lysis buffer. HA-tagged proteins were eluted in 60 µL pH 2.0 buffer (Pierce) for 10 min at 50 °C to dissociate proteins from the antibody-conjugated beads. The elutions were immediately neutralized 1:10 with pH 8.5 neutralization buffer (Pierce).

### Mass spectrometry sample preparation

45 µL of each IP elution was combined with 15 µL of 4X Laemmli sample buffer supplemented with BME (BioRad), boiled for 10 min at 95 °C, and loaded on a Bolt 4-12% Bis-Tris gel (Invitrogen). The samples were resolved for approximately 8 min at 150V. The gel was washed once in UltraPure water (Thermo), fixed in 50% methanol and 7% acetic acid for 15 min, followed by 3 additional washes with UltraPure water. The gel was stained for 10 min with GelCode Blue (Thermo) and washed with UltraPure water for an additional 20 min. One gel band (approx. 1.5 cm in size) for each condition was excised and de-stained for 2 hours in a 50% methanol and 10% acetic acid solution, followed by a 30 min soak in UltraPure water. Each gel slice was cut into 1 mm x 1 mm squares, covered in 1% acetic acid solution, and stored at 4 °C until the in-gel digestion could be performed.

To prepare samples for mass spectrometry, the 1% acetic acid solution was removed, 10 µl of 50 mM DTT was added, and volume was increased to 100 µl with 50 mM ammonium bicarbonate. Samples were incubated at 55 °C for 30 min. Samples were then brought down to RT, DTT solution was removed, 10 µl of 100 mM acrylamide (propionamide) was added and volume was again normalized to 100 µl with 50 mM ammonium bicarbonate followed by an incubation at RT for 30 min. Acrylamide solution was removed, 10 µl (0.125 µg) of Trypsin/LysC (Promega) solution was added and another 50 µl of 50 mM ammonium bicarbonate was added to cover the gel pieces. Samples were incubated overnight at 37 °C for peptide digestion. Solution consisting of digested peptides was collected in fresh Eppendorf tubes, 50 µl of extraction buffer (70% acetonitrile, 29% water, 1% formic acid) was added to gel pieces, incubated at 37 °C for 10 min, centrifuged at 10,000 x g for 2 minutes and collected in the same tubes consisting of previous elute. This extraction was repeated one more time. Collected extracted peptides were dried to completion in a speed vac and stored at 4 °C until ready for mass spectrometry.

### Mass spectrometry

Eluted, dried peptides were dissolved in 12.5 μl of 2% acetonitrile and 0.1% formic acid and 3 μl was injected into an in-house packed C18 reversed phase analytical column (15 cm in length). Peptides were separated using a Waters M-Class UPLC, operated at 450 nL/min using a linear 80 minute gradient from 4% mobile phase B to 40% B. Mobile phase A consisted of 0.2% formic acid, 99.8% water, Mobile phase B was 0.2% formic acid, 99.8% acetonitrile. Ions were detected using an Orbitrap Fusion mass spectrometer operating in a data dependent fashion using typical “top speed” methodologies. Ions were selected for fragmentation based on the most intense multiply charged precursor ions using Collision induced dissociation (CID). Data from these analyses was then transferred for analysis.

### Mass spectrometric analysis

The .RAW data were searched using MaxQuant version 1.6.1.0 (36) against the canonical human database from UniProt, *Toxoplasma* GT1 databases from ToxoDB (versions 7.3 and 37.0), and the built-in contaminant database. Specific parameters used in the MaxQuant analysis can be found in Supplemental Table 3. Peptide and protein identifications were filtered to a 1% false discovery rate (FDR) and reversed proteins, contaminants, and proteins only identified by a single modification site, were removed from the dataset. HCE1HA enrichment over the non-HA tagged RH was determined by adding 1 to each spectral count (MS/MS count), calculating the relative spectral counts observed in each experiment (MS/MS counts for each protein / total MS/MS counts for all proteins in that experiment), and then calculating the relative spectral counts observed for each protein in the two samples.

### Statistical Analyses

Statistical analysis was performed with Prism version 8 software. Analysis of plaque size was performed using Student’s t test.

## Acknowledgments

We thank all members of our laboratory and Drs. Julien Sage, Jan Skotheim, Seth Rubin and Joe Lipsick for helpful comments, as well as Melanie Espiritu for help with tissue culture and ordering. We also thank Jeff Huleatt for help with data analysis and Drew Etheridge for interesting discussion and exchange of data prior to publication.

Special thanks goes to the Vincent Coates Foundation Mass Spectrometry Laboratory, Stanford University Mass Spectrometry (SUMS) for assistance in processing mass spectrometry samples.

## Author Contributions

MWP, AN, AMC were all active in experimental design, data collection and interpretation of the data. MWP and JCB were jointly responsible for project conception, design, analysis of the data, and drafting the manuscript. All authors have approved the submitted version and have agreed to be responsible for the data presented here.

---

**Supplemental Figure 1.**
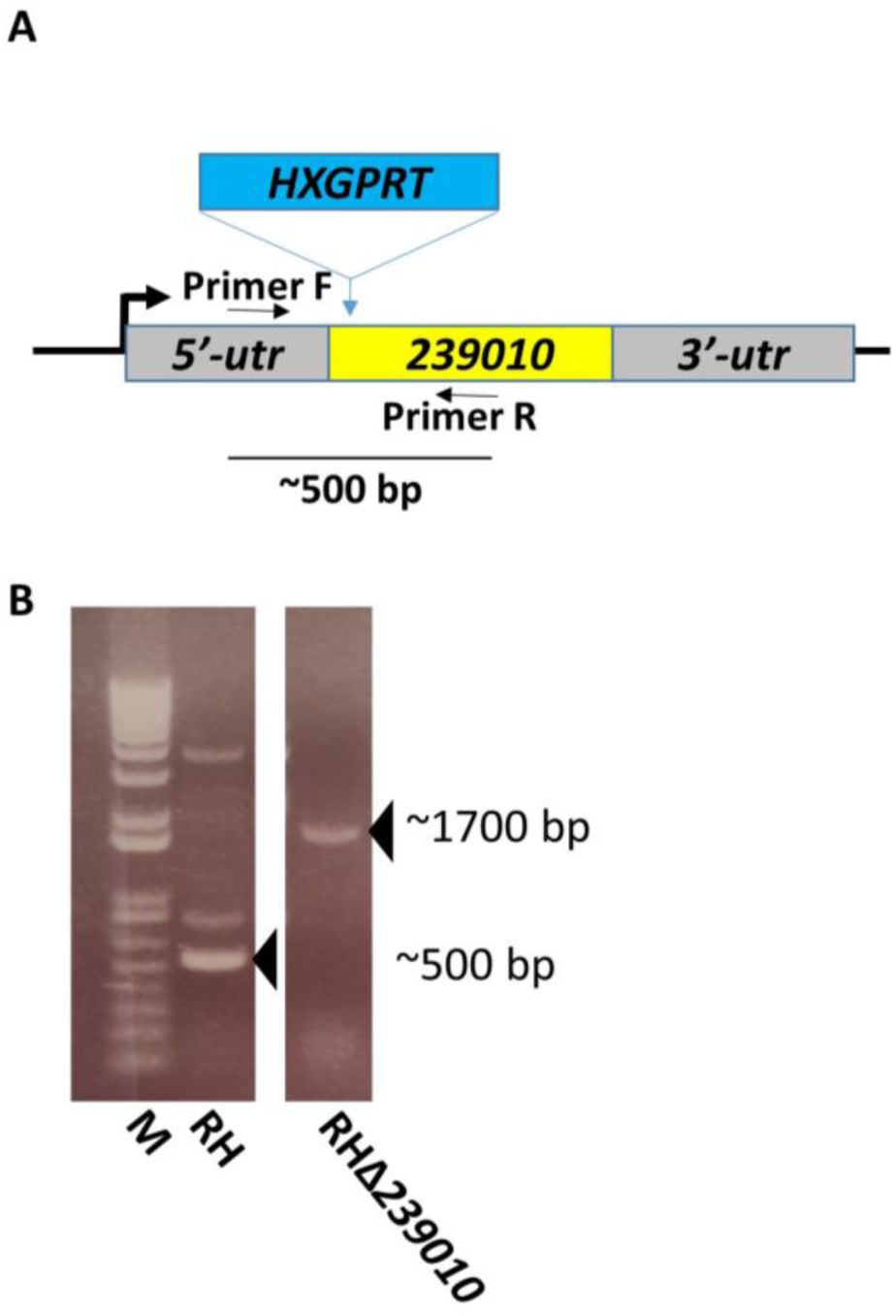
Deletion of the single exon gene *TGGT1_239010* by CRISPR-targeted insertion of *HXGPRT*. **A.** Schematic of CRISPR-mediated disruption of *TGGT1_239010* followed by insertion of the *HXGPRT* gene for selection. Primers flanking the guide-targeted region, indicated by Forward and Reverse, were constructed to amplify a ~500 base pair region of the gene. **B**. PCR amplification of DNA from wild-type RH and a HXGPRT^+^ strain with disruption of the *TGGT1_239010* (RH*Δ239010*) using the forward and reverse primers shown in A. The size markers (M) reveal bands of the expected size in the RH strain (~500 bp) and in the RH*Δ239010* strain (~1700 bp). Sequencing of the amplification products confirmed the identities of these bands as the result of PCR, as predicted from the schematic in A.

**Supplemental Figure 2.**
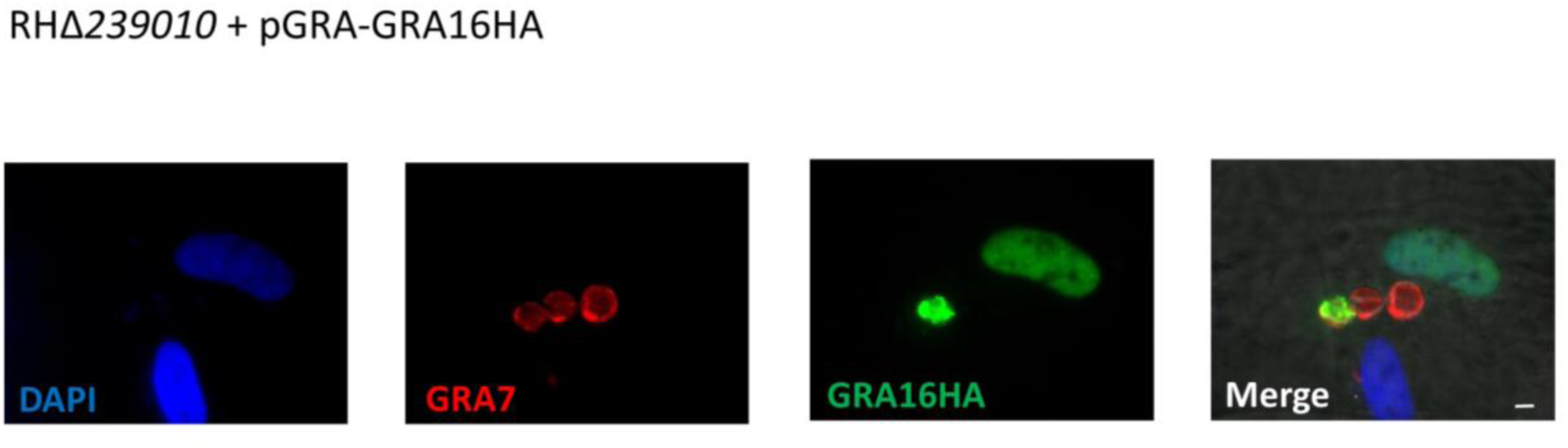
Disruption of *TGGT1_239010* does not prevent GRA16 from exiting the parasitophorous vacuole and accessing the host cell nucleus. RHΔ*239010* was transiently transfected with the plasmid pGRA-GRA16HA, and GRA16HA localization (green) was assessed by IFA. GRA7 (red) was used to visualize the parasitophorous vacuoles, whether they harbored transfected parasites or not. White scale bar is 5 μm.

**Supplemental Figure 3.**
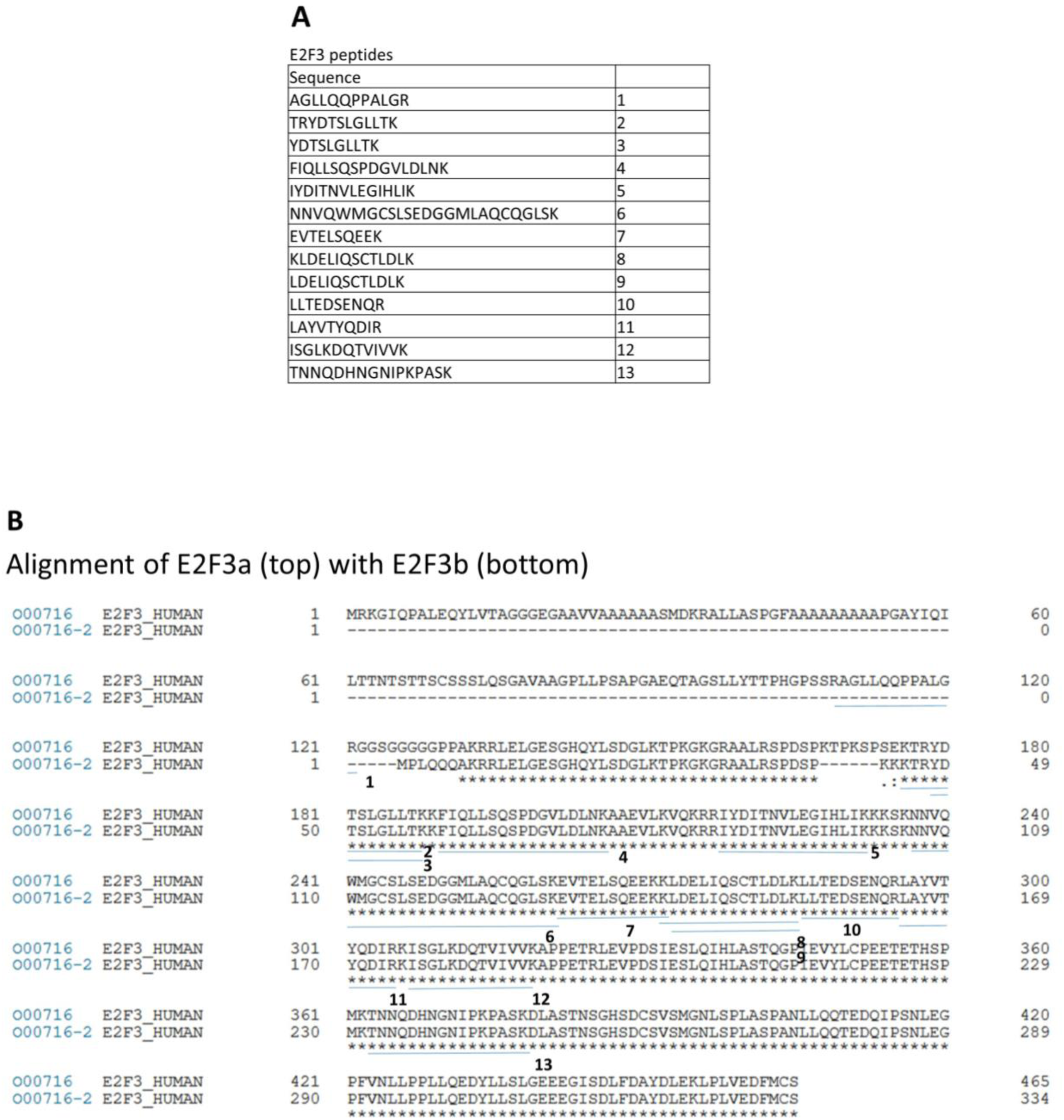
Alignment of E2F3a and E2F3b and identification of peptide fragments that immunoprecipitated with HCE1. Peptide fragments 1-13 are displayed in a table (**A**), and these were mapped to the alignment of E2F3a and E2F3b (**B**). Blue lines represent the peptide fragment and the fragment number is immediately to the right of the blue line. Only peptide fragment 1, for which there was one spectral count found to associate with HCE1 in each IP, mapped to a region unique to E2F3a. All other peptide fragments mapped to regions common to E2F3a and E2F3b.

**Supplemental Table 1.** RNASeq results for the 602 annotated genes shown in Figure 3 that are expressed in RH-infected HFF at least 2.5-fold higher than they are in RHΔ*239010*-infected HFF. RPKM values are given for each gene during mock infection, infection with RH, RHΔ*239010*, and RHΔ*myr1*. Genes are ranked in decreasing order of the ratio of expression in RH/RHΔ*239010*. For instances where there was no expression of the gene in RHΔ*239010*-infected cells, genes with highest expression in RH-infected cells are listed first.

**Supplemental Table 2.** A list of the host genes that were upregulated in RH-infected HFFs but not those infected with RHΔ*239010* parasites and that contributed to the three GSEA gene sets that were significantly enriched in the RH-vs. RHΔ*239010*-infected cells, as shown in Figure 3.

**Supplemental Table 3.** Mass spectrometry analysis parameters and results for proteins that co-immunoprecipitate with HCE1HA-expressing and untagged RH parasites. For all sheets, the majority protein IDs, i.e. the proteins which contain at least half of the peptides belonging to a protein group (grouping of proteins which cannot be unambiguously identified by unique peptides), and the number of spectral counts (MS/MS count) corresponding to each grouping are shown. The bait protein, HCE1, is highlighted in yellow. Protein IDs associated with human proteins are colored in blue. The gene product (for *Toxoplasma* proteins) or associated gene name (for human proteins) for the first listed protein ID in each row is shown in the Description column. Sheet 1 (“All_proteins”) shows the experimental data sets for all proteins, both human and *Toxoplasma*, listed in rank order by enrichment score (HCE1/RH, as further elaborated in the Materials & Methods) in IP #1. Sheet 2 (“Human_proteins”) shows the experimental data sets for human proteins only listed in rank order by enrichment score in IP #1. Sheet 3 (“Toxo_proteins”) shows the experimental data for *Toxoplasma* proteins only listed in rank order by enrichment score in IP #1. Sheet 4 (“Parameters”) shows the parameters used in the MaxQuant analysis.

